# Strategies of developmental vestibular plasticity after unilateral embryonic ear removal in *Xenopus laevis*

**DOI:** 10.1101/2022.06.08.495274

**Authors:** Clayton Gordy, Hans Straka

**Affiliations:** Faculty of Biology, Ludwig-Maximilians-University Munich, Großhaderner Str. 2, 82152 Planegg, Germany; Graduate School of Systemic Neurosciences, Ludwig-Maximilians-University Munich, Großhaderner Str. 2, 82152 Planegg, Germany

**Keywords:** inner ear, vestibular, plasticity, *Xenopus laevis*, brainstem, vestibulo-ocular reflex, development, eye movements, extraocular motoneurons

## Abstract

Appropriate gaze control mechanisms rely on bilateral mirror-symmetric vestibular endorgans, central circuits, and extraocular motor effectors. Embryonic removal of one inner ear prior to the formation of these structures was used to evaluate the extent to which animals can develop appropriate motor outputs in the presence of only a singular inner ear. Near-congenital one-eared tadpoles subjected to separate or combinatorial visuo-vestibular motion stimulation exhibited vestibular-ocular reflexes, though smaller in gain compared to controls, whereas isolated visuo-motor responses remained largely unaltered. Surprisingly, direction-specific responses in one-eared tadpoles revealed a spectrum of vestibulo-ocular reflex performance, where in many cases rather robust responses were observed during head motion towards the missing ear. Rotation toward the remaining ear elicited eye movements which were often severely attenuated. This suggests that embryonically generated one-eared animals develop bidirectional vestibular signal detection and downstream processing capacities that allow activation of robust compensatory eye movements. However, this capability seems to occur at the expense of performance of vestibular reflexes during motion in the canonical activation direction of the singular ear. Consequently, the development of central vestibulo-motor circuits in one-eared animals likely relies on strategies in which the outcome for gaze stabilization is driven by balanced homeostatic levels and individual temporal tuning of abducens motoneuron activity. Despite the lack of bilateral symmetric vestibular afferent input, the developing central nervous system is capable of creating a framework for symmetric sensorimotor transformations.

## Introduction

Head movements are detected and mechano-electrically transduced into neuronal signals by vestibular organs in the inner ear (Angelaki and Cullen, 2008; Dieterich and Brandt, 2015). Following vectorial decomposition by semicircular canal and otolith organs, bilateral signals are reconstructed through spatially-specific integration in central circuits and contribute to behaviors which stabilize posture and gaze during active and passive movements (Straka and Gordy, 2020). A key feature of this computation is the mirror-symmetric arrangement of sensory epithelia (Fritzsch and Straka, 2014) and the interconnection of the vestibular nuclei across the midline by commissural pathways (Malinvaud et al., 2010). Despite this bilaterality, such mirror-symmetry generates motion sensors on both sides that are largely distinct from each other with respect to directional preference. Bilateral vestibular organs therefore represent complementary structures with distinct sensitivity domains rather than simple duplications with redundant functionality (Chagnaud et al., 2017). Thus, encoding and representation of multi-dimensional head/body movements depends on the morpho-physiological integrity of vestibular sensors within the two inner ears.

Disruption of bilateral processing, such as during an acute unilateral loss of inner ear function or inappropriate peripheral signaling, results in an impairment of self-motion encoding due to insufficient and asymmetric information from symmetric sensors. Immediate behavioral effects include dizziness, vertigo, spontaneous nystagmus, and deterioration of orientation and navigational skills (Zhao et al., 2008; see Fetter, 2016). These pathological reactions derive from excessive bilateral asymmetric activity of central vestibular circuits combined with the subsequent failure to produce adequate gaze- and posture-stabilizing neuronal commands. In addition, asymmetric neuronal activity is centrally represented as being in mismatch with other motion-related sensory signals such as visual image motion or limb/neck proprioceptive inputs (see e.g., Curthoys, 2000; Dutia, 2010; Strupp and Brandt, 2013). However, these impairments abate, at least partially, over time due to plasticity processes in bilateral central circuits which are distributed across various regions of the central nervous system (CNS), and occur at molecular, cellular, and anatomical levels, which collectively permit readjustments in computational strategies to alleviate the consequences of peripheral imbalance (Llinás and Walton, 1979; Dieringer, 1995; Straka et al., 2005).

The remarkable plasticity of vestibular signal processing after a unilateral vestibular loss has been extensively used to study the principles of “vestibular compensation” following a variety of protocols (e.g., Curthoys, 2000). These studies were usually conducted in adult (e.g., Dieringer, 1995) or juvenile (Lambert et al., 2013) vertebrates with a functional vestibular sensory periphery and central pathways. In this manner, unilateral impairments of inner ear function induced a loss of signal processing in already well-established, entrained, and spatio-temporally tuned circuits. Under such circumstances, vestibular lesion-induced plasticity must cope with preexisting bilateral symmetric circuits and resultant computations. In contrast, unilateral excision of the embryonic otic placode, which develops into all sensory and non-sensory tissues of the inner ear, prior to the formation of central pathways (Elliott and Fritzsch, 2010; Elliott et al., 2015a, b) might reveal plasticity processes that permit vestibular circuits to develop and function based on sensory inputs only from a single inner ear into circuits which have only ever received such unilateral input. This generates a developmental condition where relevant brainstem vestibular circuits control bilateral gaze- and posture-stabilizing motor elements from unilateral vestibular inputs alone.

Here, we demonstrate that unilateral embryonic removal of the otic placode causes one-eared tadpoles to exhibit a remarkable degree of developmental vestibular plasticity. These tadpoles develop without morphological signs typical for a unilateral vestibular loss. Gaze-stabilizing vestibulo-motor responses exhibit appropriate spatiotemporal dynamics during bidirectional motion stimulation. Behavioral analyses during unidirectional motion, as well as electrophysiological evidence, suggest that central circuits have adapted to respond to oscillatory head motion within the singular ear, with minimal contributions by motion-sensitive visual pathways. Collectively these results highlight the ability of the central nervous system to develop appropriate motion direction-specific gaze-stabilizing behaviors following ontogenetic assembly of circuits in the absence of bilateral signaling.

## Results

### Vestibular-evoked eye movements in one-eared tadpoles

One-eared tadpoles were generated by unilateral removal of the left otic placode at embryonic stages 25-27 (Figure 1A, left; Supplemental video 1). Removal of the otic placode at these developmental stages has previously been demonstrated in *Xenopus laevis* to selectively and completely remove inner ear endorgans and corresponding neurosensory elements (Fritzsch, 1990; Elliott and Fritzsch, 2010). The absence of the entire ear and its resulting lack of peripheral sensory components was confirmed beginning at stage 46 (Figure 1A), a developmental period where the high transparency of *Xenopus* tadpoles allows direct visual assessment of the presence of inner ear structures (Supplemental Figure 1A_1_-A_2_, C_1_-C_2_). As expected, stage 46 one-eared animals lacked recognizable inner ear gross-histological structures (Supplemental figure 1B_1_, D_1,_ asterisk) as well as neurosensory elements such as hair cells and associated vestibular afferent fibers (Figure 1A, extirpated side; Supplemental figure 1B_2_-B_3_, D_2_-D_3_). Using myosin-VI and acetylated-tubulin as selective markers for hair cells and nerve fibers, respectively, a clear absence of innervated sensory epithelia on the operated side, compared to the unmanipulated side was revealed (Figure 1A, Supplemental figure 1D_2_), confirming the successful and reliable embryonic removal of one ear.

**Figure 1.**
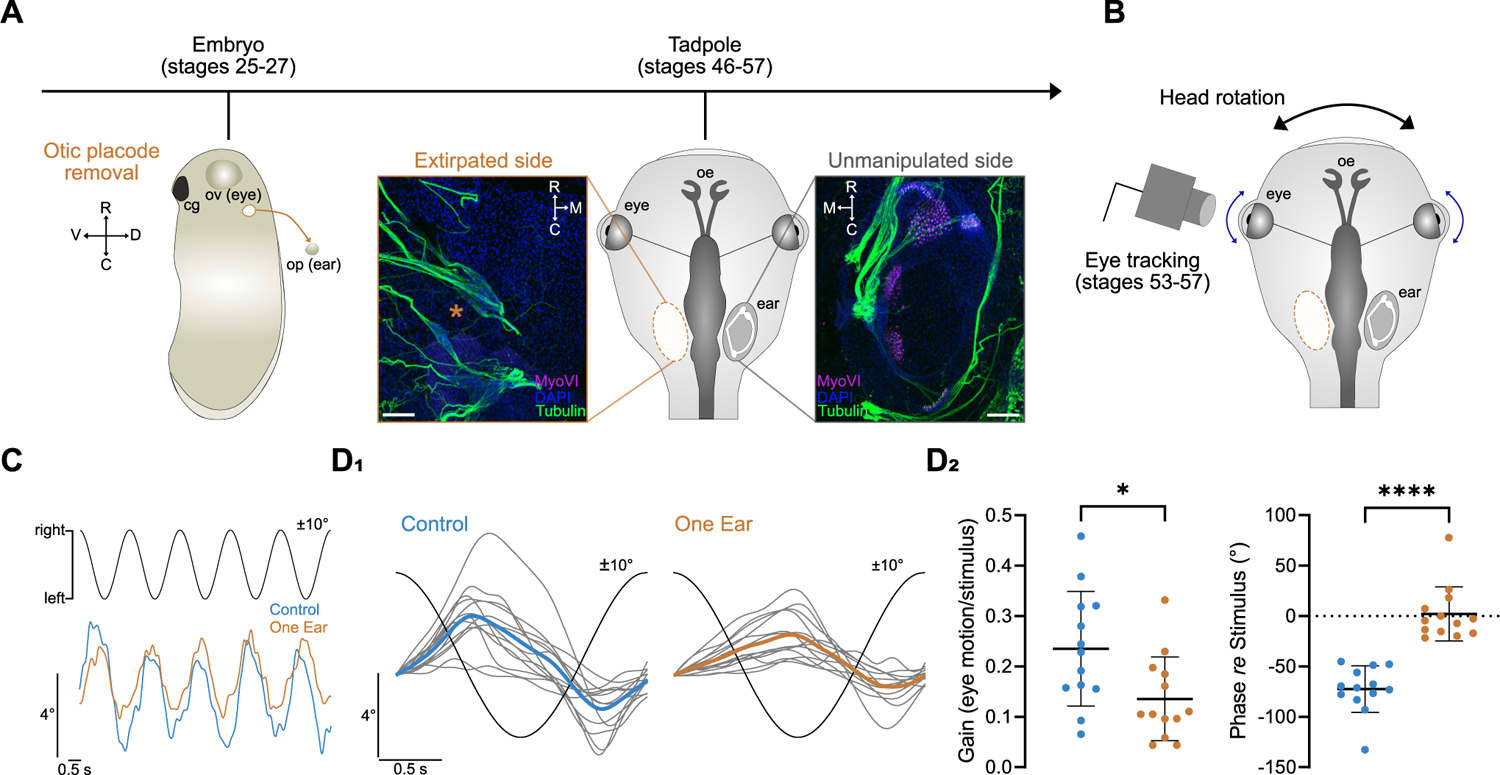
Vestibulo-ocular reflex performance in one-eared *Xenopus laevis* tadpoles. **(A)** Schematic depicting the experimental procedure and developmental timeline following unilateral embryonic removal of the otic placode (stages 25-27; lateral view) followed by rearing of the one-eared embryos to tadpole stages (stage 46-57; dorsal view); note the lack of the left inner ear (orange *) and corresponding neurosensory and accessory otic structures, illustrated by images from the left (Extirpated side) and right side (Unmanipulated side) of a stage 46 larva, with whole-mount antibody stainings against neurons (acetylated tubulin, green) and hair cells (myosin-VI, red) in the otic region. **(B)** Schematic of a semi-intact preparation used for functional profiling of control and one-eared tadpoles during horizontal sinusoidal rotation coupled with live motion-tracking of both eyes. **(C)** Representative example of oppositely-directed, compensatory eye oscillations (lower traces) during five cycles of horizontal sinusoidal head rotation (±10°, peak velocity ±31.4°/s) at 0.5 Hz (upper trace) in an unmanipulated control (blue) and a one-eared (orange) tadpole. Responses are averages of both eyes, respectively. (**D**) Averaged responses over a single horizontal rotation cycle of controls (*n* = 13, individual gray traces; from 6-40 cycles) and one-eared animals (*n* = 13, individual gray traces; from 12-66 cycles); blue and orange traces represent the population mean response over one motion cycle (black trace) for the respective group of animals (**D_1_**); averaged responses were used to individually calculate the gain (left in **D_2_**) and phase value *re* stimulus position (right in **D_2_**). Significance levels are indicated by asterisks: * *p* ≤ 0.05, **** *p* ≤ 0.0001 (Mann-Whitney *U*-test). R, rostral; C, caudal; V, ventral; D, dorsal; M, medial; op, otic placode; ov, optic vesicle; cg, cement gland; oe, olfactory epithelium. Immunohistochemical stainings in **A** were counterstained with the nuclear marker DAPI. Scale bars in **A** are 100 µm.

Given that semicircular canals in *Xenopus laevis* tadpoles become functional at stage 48 (Lambert et al., 2008) and only elicit robust angular vestibulo-ocular reflexes (aVOR) after having reached stage 52/53, one-eared tadpoles were reared to this developmental stage (Figure 1B, Supplemental figure 1E_1_-F_2_). *In vitro* preparations of unmanipulated two-eared controls and one-eared tadpoles (Supplemental figure 1E_1_-E_2_, F_1_-F_2_) were used to assess the performance of gaze stabilizing vestibulo-ocular motor responses by eye motion tracking during horizontal rotation on a motion platform in complete darkness (Figure 1B). Sinusoidal rotation of unmanipulated control tadpoles in the dark at 0.5 Hz with a peak velocity of ±31.4°/s, corresponding to positional excursions of ±10° (Figure 1C, top trace), evoked vestibular-driven compensatory eye movements (without intermittent fast-phases) that were positional stimulus-timed and oppositely directed, features that are characteristic for the aVOR in *Xenopus* tadpoles (Figure 1C, bottom traces). Across all control animals, an average over single cycles (6-40 cycles) in the dark (Figure 1D_1_, left) exhibited a response gain (eye motion amplitude / stimulus position amplitude) of 0.24 ±0.11 (Figure 1D_2_, left; mean ±SD, *n* = 13) and a considerable phase-lead *re* stimulus position of −72.46° ±23.13° (Figure 1D_2_, right; mean ±SD, *n* = 13).

Despite the complete absence of inner ear endorgans on the left side, one-eared animals subjected to the same stimulation paradigm also exhibited oppositely directed eye movements indicative of a functional aVOR (Figure 1D_1_, right). Response magnitudes, obtained by averaging over multiple cycles (12-66 cycles) presented with gain values of 0.14 ±0.08 (Figure 1D_2_, left, mean ±SD, *n* = 13) and a response peak that was approximately in phase *re* stimulus position (2.08° ±26.90°; Figure 1D_2_, right, mean ±SD, *n* = 13). Statistical comparison of eye movements between one-eared tadpoles and controls revealed a significant reduction of the response gain (Figure 1D_2_, left, *p* = 0.0338; Mann-Whitney *U*-test). In addition, the pronounced phase-lead of the peak responses relative to stimulus position in darkness was significantly delayed in one-eared tadpoles relative to controls (Figure 1D_2_, right, *p* < 0.0001; Mann-Whitney *U*-test). Accordingly, these data demonstrate that one-eared tadpoles are able to execute a horizontal aVOR in darkness despite the lack of bilateral mirror-symmetric endorgans and indicate that the remaining intact inner ear is sufficient to produce gaze-stabilizing extraocular motor commands, even though with reduced efficacy. The phase-relationship of the responses in one-eared animals suggests a considerable temporal delay in the processing of signals from the right, singular, inner ear.

### Directional contributions of singular ears during horizontal aVOR

Head rotation is normally encoded by direction-specific strengthening/attenuation of vestibular nerve afferent signals (Paulin and Hoffman, 2019). In one-eared tadpoles, which maintain the ability to encode oscillatory motion in darkness (Figure 1), a single semicircular canal was found to be sufficient for eliciting a bidirectional horizontal aVOR. However, to separately investigate the directional contributions of a singular ear to leftward *versus* rightward head movements, eye motion amplitudes were evaluated over the first half-cycle of stimulation bouts during platform rotation exclusively to the left or to the right (Figure 2A). Eye movements during these half-cycle periods would therefore derive only from a unidirectional motion away from the singular ear (contraversive) or toward this intact ear (ipsiversive). In two-eared unmanipulated controls, eye movements evoked by unidirectional motion in the dark towards the left (Figure 2B, left; blue traces) or the right (Figure 2B, right; blue traces) were predictably opposite and statistically no different in response strength to stimulus direction within individual animals with mean amplitudes of 5.96° ±1.44° and 5.93° ±1.63°, respectively (Figure 2C, left, mean ±SD, *p* > 0.9999; Wilcoxon signed-rank test, *n* = 4 pairs). Eye movements in one-eared tadpoles evoked by leftward, contraversive, motion in the dark surprisingly were rather variable but astonishingly also robust and in opposition to head movements (Figure 2B, left; orange traces). Rightward ipsiversive motion, i.e., towards the side of the intact, singular ear, evoked responses that were even more variable between different animals, both in direction and magnitude (Figure 2B, right; orange traces). In addition, these eye movements were generally smaller than those driven by contraversive motion with mean amplitudes of 1.43° ±2.05° and 3.82° ±1.88°, respectively (Figure 2C, right, mean ±SD, *p* = 0.0420, Wilcoxon signed-rank test, *n* = 11 pairs). Surprisingly, despite the inner ear being intact on the right side, aVOR responses elicited by a rightward ipsiversive motion in one-eared tadpoles were severely impaired compared to controls (Figure 2D, ipsi, *p* = 0.0002; Mann-Whitney *U*-test). Such a significant impairment was also found for contraversive motion-driven eye movements toward the side lacking an ear (Figure 2D, contra, *p* = 0.0136; Mann-Whitney *U*-test), although this outcome was expectable given the lack of canonical sensitivity for motion toward the impaired inner ear (see e.g., Soupiadou et al 2020). Thus, these sets of data indicate that one-eared tadpoles developed bidirectional vestibular detection and signal processing capacities that allow activating compensatory eye movements during rotation towards the side lacking an ear. However, this directional contribution obviously occurs at the expense of the performance of the aVOR towards the intact side, which becomes profoundly compromised during this process (Figure 2D).

**Figure 2.**
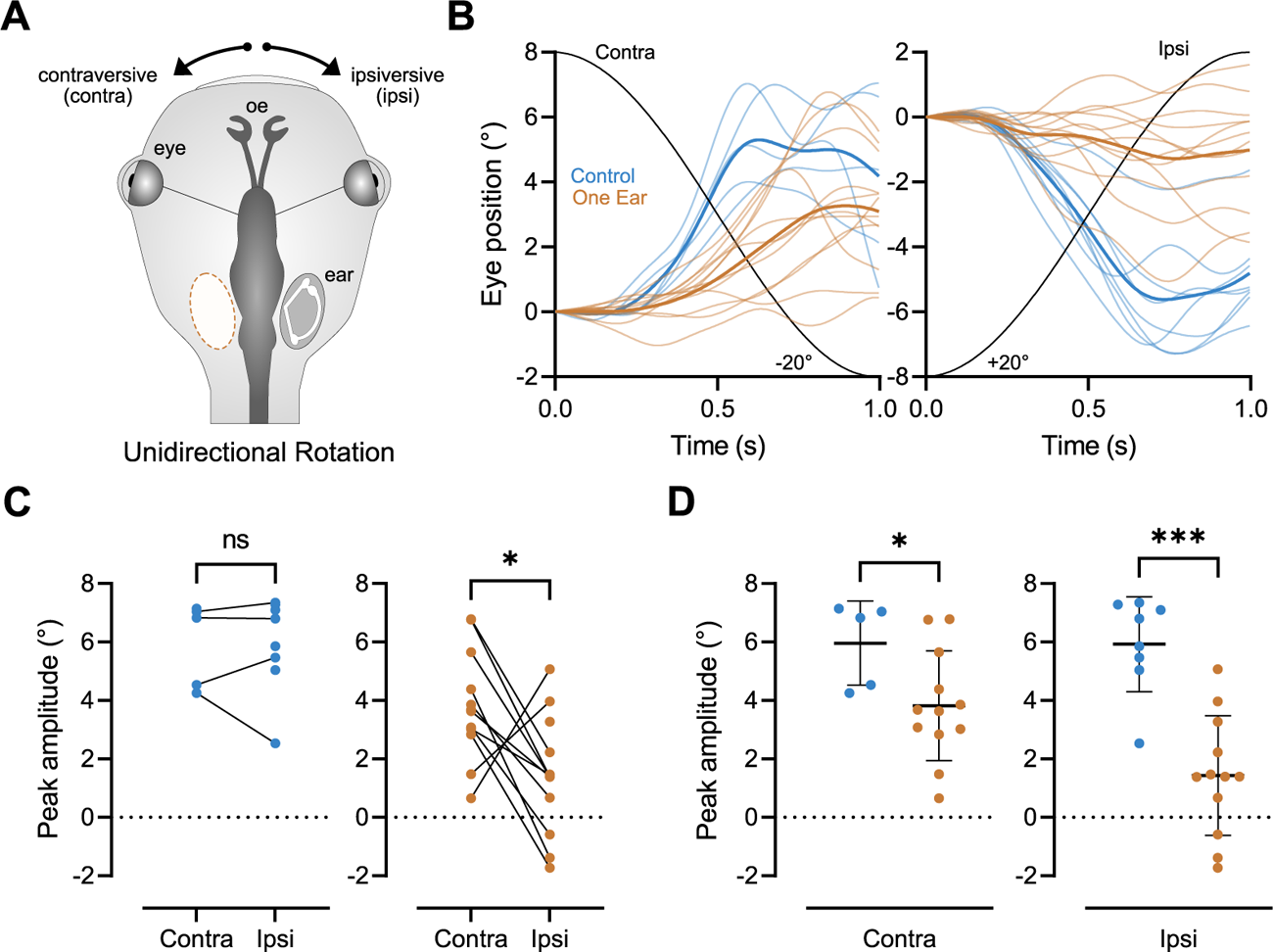
Directional sensitivities of singular ears during horizontal aVOR. **(A)** Schematic depicting unidirectional horizontal angular rotation of control and one-eared animals; rotations were performed either toward (ipsiversive, ipsi) or away from the residual singular ear (contraversive, contra) without oscillation between the two directions. **(B)** Eye movements of individual control (*n* = 9; thin blue traces) and one-eared (*n* = 13; thin orange traces) animals during unidirectional rotation, averaged over 1-6 half-cycles, respectively, that were obtained from the onset of sinusoidal stimulus events shown in Figure 1D; thick blue and orange traces represent respective population means. **(C, D)** Comparison of peak response amplitudes during contraversive and ipsiversive positional excursions within **(C)** controls (blue) and one-eared animals (orange), respectively, and for the two directions between controls and one-eared animals **(D)**; dotted lines in **C** and **D** represent the reversal lines of eye motion direction; note that peak amplitudes during ipsiversive rotations were inverted to facilitate a comparison between the responses for the two stimulus directions; significance levels are indicated by asterisks: * *p* ≤ 0.05 (Wilcoxon signed-rank test) *** *p* ≤ 0.001 (Mann-Whitney *U*-test).

### Visuo-vestibular plasticity and influence on gaze-stabilizing reflexes

Motion-related sensory signals are known to participate in plasticity processes aiding recovery of acute vestibular loss (e.g., Smith, 2022). In aquatic organisms, visual scene motion is a significant contributor in neuronal computations of self-motion behaviors (Roeser and Baier, 2003), particularly through optokinetic reflex (OKR) circuits which operate synergistically with aVOR signals (Souipadou et al., 2020). To investigate the extent that visual image motion assists vestibular-evoked eye movements in one-eared animals, tadpoles were subjected to horizontal sinusoidal rotation of the platform in the presence of a world-stationary illuminated black and white-striped visual pattern (light; Figure 3A). This experimental approach caused a synergistic activation of a horizontal aVOR and an OKR. Eye movements evoked in un-manipulated control tadpoles under this condition were oppositely directed (Figure 3B) with gain magnitudes of 0.22 ± 0.08 (Figure 3C_1_, mean ±SD, *n* = 13) and were timed with stimulus position with relatively small phase leads of −18.74° ± 17.97° *re* head position (Figure 3D_1_, mean ±SD, *n* = 13). When compared to similarly evoked movements in darkness (dark; Figure 1, Figure 3B, dotted blue line) gain magnitudes were found to be no different to eye movements evoked in light (Figure 3C_1_, *p* = 0.6355; Wilcoxon signed-rank test, *n* = 13 pairs). In contrast, quantification of phase relationships revealed that eye movements evoked in light were considerably more in phase with stimulus head position (Figure 3D_1_, *p* = 0.0002; Wilcoxon signed-rank test, *n* = 13 pairs). Such a relationship between aVOR responses in the presence of a world-stationary visual scene and aVOR in darkness in two-eared control animals complies with the canonical impact of concurrent visual motion signals on gaze-stabilizing VOR behaviors, where visual image motion serves as ongoing feed-back to adjust the VOR dynamically with only minor influences on response magnitude (see Straka and Dieringer, 2004). In one-eared tadpoles, vestibular-evoked eye movements in light were stimulus-timed and oppositely directed with gain and phase magnitudes of 0.16 ± 0.08 and 4.58° ± 14.71°, respectively (Figure 3C_2_, D_2_, mean ±SD, *n* = 13). Similar to un-manipulated controls, gain magnitudes did not differ statistically between light and dark conditions (Figure 3C_2_, *p* = 0.1272; Wilcoxon signed-rank test, *n* = 13 pairs). However, in contrast, vestibular-evoked eye movements in one-eared animals in light did not exhibit a further in-phase shift relative to head position as observed in control animals (Figure 3D_2_, *p* = 0.3396; Wilcoxon signed-rank test, *n* = 13). This suggests the lack of a behaviorally observable influence of visual image motion on aVOR circuits in these animals.

**Figure 3.**
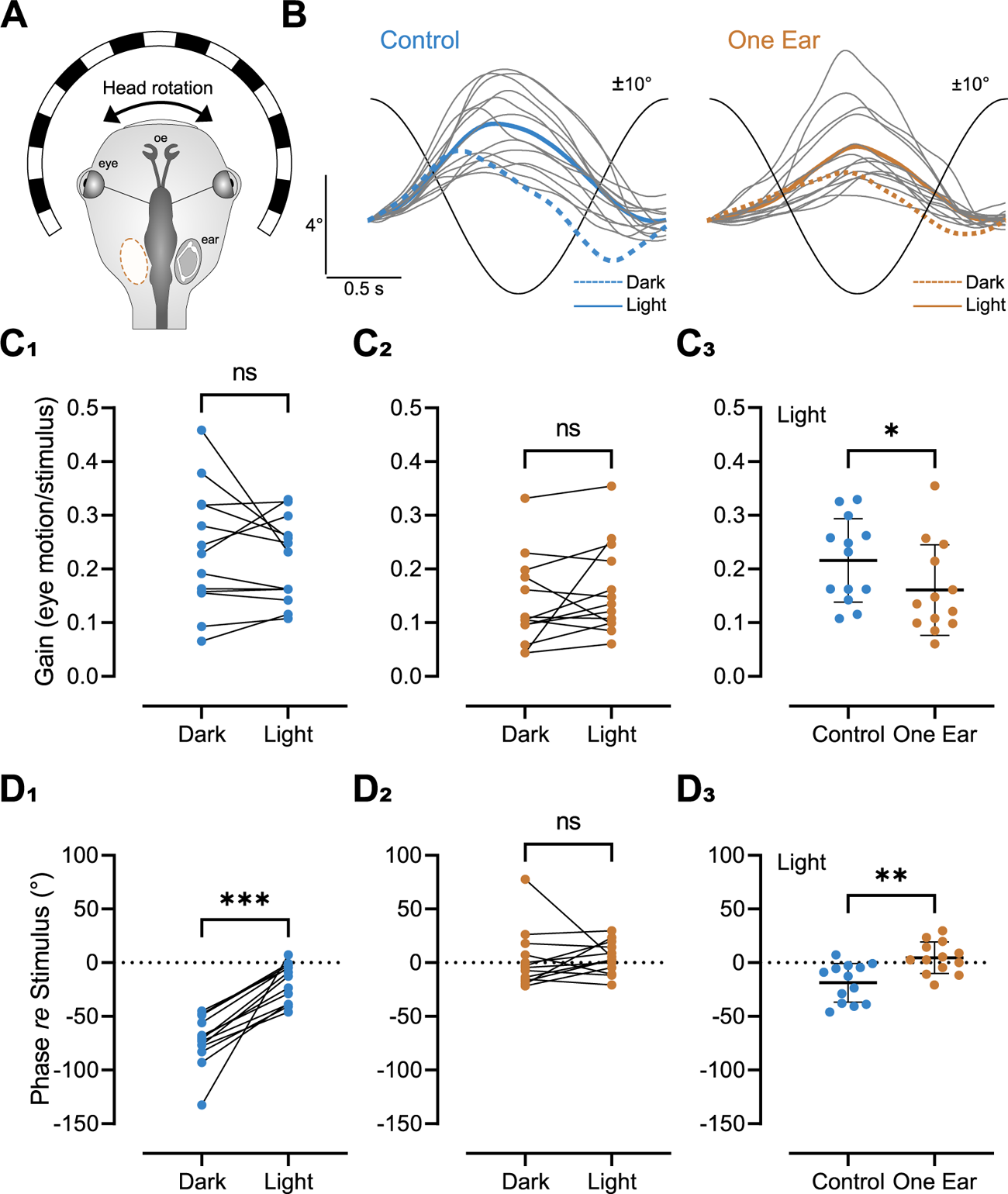
Visuo-vestibular reflex plasticity. **(A)** Schematic depicting the experimental condition that consisted of a horizontal sinusoidal head rotation in the presence of a world-stationary, illuminated black and white-striped visual pattern (Light). **(B)** Averaged responses over a single head motion cycle in light (gray traces from 6-77 cycles, respectively) and population means (solid-colored traces) in controls (*n* = 13) and one-eared animals (*n* = 13); dotted blue and orange traces depict population means obtained from head rotations in darkness (Dark) illustrated in Figure 1D; black sine waves indicate the stimulus position. (**C-D**) Gain **(C_1_-C_3_)** and phase *re* stimulus position **(D_1_-D_3_)** calculated from averaged responses over a single motion cycle in Dark and Light conditions of controls **(C_1,_ D_1_)** and one-eared animals **(C_2,_ D_2_)**; respective values for the light condition in the two experimental groups are compared in **C_3,_ D_3_**. Significance levels are indicated by asterisks: * *p* ≤ 0.05, ** *p* ≤ 0.01, *** *p* ≤ 0.001 (Wilcoxon signed-rank test in **D_1_**, Mann-Whitney *U*-test in **C_3_, D_3_**). Horizontal dotted lines in **D_1-3_** at 0° indicate phase alignment of the stimulus.

Comparison of the performance of stimulus-evoked eye motion in one-eared animals and unmanipulated controls in the presence of an illuminated visual pattern revealed smaller overall gain values as well as significantly more in phase responses for one-eared animals (Figure 3C_3_, D_3_, *p* = 0.0441; Mann-Whitney *U*-test and *p* = 0.0020; Mann-Whitney *U*-test, respectively). This is consistent with differences observed in darkness (Figure 1D_2_) and indicates that the canonical influence of visual scene motion on temporal adjustments of the VOR does not occur in one-eared animals. Furthermore, these animals continue to perform statistically less robust than controls even in the presence of a visual scene.

To rule out that the optokinetic circuit itself was not disrupted as a result of the embryonic loss of one ear, separate activation by sinusoidal motion of a vertically-striped black and white pattern at different frequencies (0.1, 0.2, 0.5 Hz, peak positional excursion of ±10°) while the head/body remained stationary was performed (Supplemental figure 2A, C). This exclusive visual scene motion provoked syndirectional eye movements with respect to the stimulus direction (Supplemental figure 2B, D). In control two-eared animals, averaged responses over single motion cycles at three different frequencies had average gains of 0.22 ±0.10, 0.11 ±0.07 and 0.05 ±0.03, respectively (mean ±SD; Supplemental Figure 2E). Comparison indicated that the response gain was statistically different between all tested frequencies, with higher frequencies evoking considerably smaller eye motion responses as expected for visual image motion processing bandwidths (Supplemental figure 2E; Friedman nonparametric test for matched pairs, *p* < 0.0001). Near similar differences were observed for phase *re* visual stimulus position relationships, with 0.5 Hz being considerably phase-lagged *re* stimulus (58.09° ±27.29°, mean ±SD) compared to the mostly in-phase responses at lower frequencies (Friedman nonparametric test for matched pairs, Dunn’s multiple comparisons test; 0.1 Hz, *p* < 0.0001; 0.2 Hz, *p* = 0.0181; Supplemental figure 2G). In one-eared tadpoles, visual motion stimulation elicited eye movements with comparable magnitudes and phase relationships, with response gains of 0.27 ±0.17, 0.15 ±0.10 and 0.05 ±0.03 (mean ±SD) for stimulus frequencies of 0.1, 0.2 and 0.5 Hz, respectively (Supplemental figure 2E). Across this frequency range, eye movements at a frequency of 0.5 Hz were considerably weaker relative to 0.1 and 0.2 Hz (Friedman nonparametric test for matched pairs, Dunn’s multiple comparisons test, *p* < 0.0001 and *p* = 0.0239, respectively). A similar relationship was found for phase characteristics of peak responses, where responses evoked at 0.5 Hz were substantially phase-lagged relative to lower frequencies (Supplemental figure 2G; 59.47° ±26.89°, mean ±SD; Friedman nonparametric test for matched pairs, Dunn’s multiple comparisons test, *p* < 0.0001 and *p* = 0.0429, respectively). Irrespective of within group differences, comparison between one- and two-eared animals revealed only very few differences in response characteristics of gain and phase (Supplemental figure 2F, H; 0.2 Hz phase comparison, *p* = 0.0355, Mann-Whitney *U*-test). Comparatively, these data demonstrate that the visuo-motor ability is overall neither impaired nor greatly enhanced in one-eared tadpoles and follows response characteristics similar to unmanipulated controls. Given the lack of additional visual image motion-mediated modulation of the aVOR in these animals (Figure 3), this collectively suggests that the vestibular circuitry and performance in one-eared tadpoles derives exclusively from sensory inputs from the remaining inner ear with little influence from visuo-motor centers.

### Physiological dynamics of one-ear-driven aVOR

To investigate the potential mechanisms used by one-eared animals to execute aVOR behaviors, electrophysiological recordings of selected extraocular motor nerves were performed. Motion of the eyes during horizontal aVOR is in part driven by the coordinated efforts of lateral recti (LR) muscles. Yoking of these muscles derives from the firing dynamics of bilateral abducens nerves which innervate each LR muscle. The discharge activity of the left (Le) and right (Ri) abducens nerve in control and one-eared animals (Figure 4A) was profiled during horizontal sinusoidal head rotation in darkness (0.5 Hz, positional excursion ±10°, peak velocity ±31.4°/s; Figure 4A).

**Figure 4.**
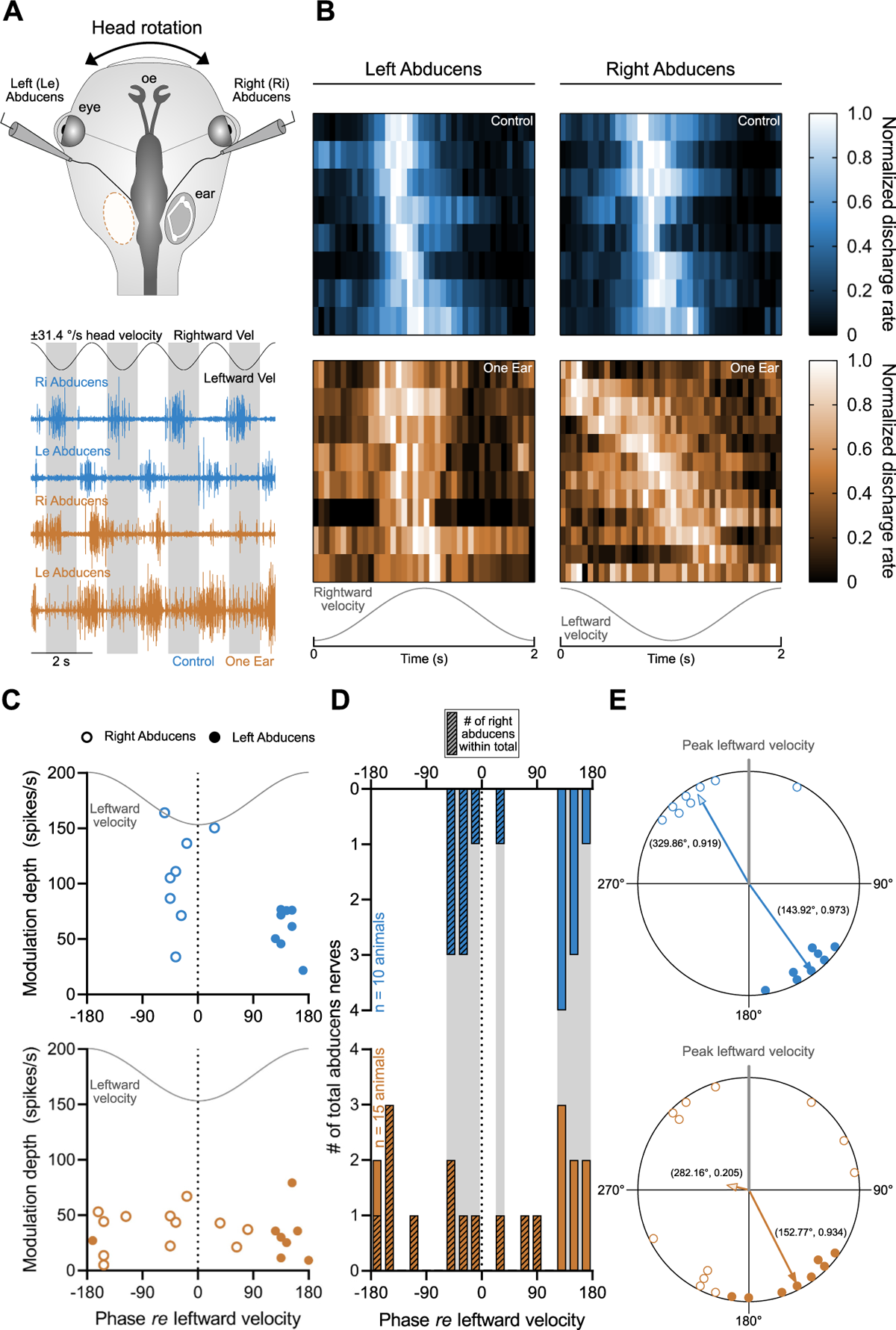
Discharge dynamics of abducens motoneurons. **(A)** Recording sites of abducens motor nerves during sinusoidal head rotation (±10° positional excursion, peak velocity of ±31.4°/s, 0.5 Hz) in darkness (upper panel); multi-unit recordings of left (Le) and right (Ri) abducens nerves (lower panel) during head rotation, corresponding to peak leftward (lower peaks) and rightward (upper peaks) velocities (Vel) of ±31.4°/s (black sinusoidal velocity trace) in two-eared control (blue) and one-eared (orange) manipulated animals; shaded regions indicate periods of leftward head motion velocity. **(B)** Heat maps visualizing peri-stimulus time histograms of normalized discharge rates over a single cycle (from 12-28 and 14-54 cycles in *n* = 10 and *n* = 15 controls and one-eared animals, respectively) during directionally specific head motion velocity (gray sinusoidal traces); horizontal heat map rows represent individual animals. **(C)** Modulation depth as a function of phase *re* peak leftward stimulus velocity for left and right abducens nerves obtained from **B**, depicting the timing of the peak discharge within the cycle; closed and open circles indicate left and right abducens nerves, respectively; note the discrete clustering of left and right abducens nerve activity in controls (upper, blue) compared to one-eared animals (lower, orange). **(D)** Frequency distribution of response phases for right and left abducens nerves, obtained from the data depicted in **C**; bar amplitudes denote the total number of nerves *per* temporal allocation; hashed bars indicate the number of right abducens nerves within the total number *per* temporal allocation. **(E)** Polar plots depicting phase deviations *re* peak leftward velocity (gray vertical line indicates phase of peak leftward velocity during stimulus motion) from **C-D** represented across 360°; arrows indicate the calculated mean vector for pooled left (filled arrowhead) and right (shaded arrowhead) abducens nerve discharge profiles in controls (upper) and one-eared (lower) animals; values next to vector arrows are respective metrics of mean angular direction and vector length (*μ, r*).

In control two-eared animals (Figure 4A, blue traces), modulation of left and right abducens nerves occurred during sinusoidal rotation in darkness in approximate phase-opposition with respect to the same-sided directional head motion velocity (see shaded gray bars). These features were consistent with the push-pull functional dynamics of extraocular motor nerves during an aVOR (Straka and Dieringer, 2004). In one-eared animals (Figure 4A, orange traces), such functional characteristics were much less consistent. While left and right abducens nerves in control animals modulated generally in phase with their opposite directional head motion velocity (Figure 4B and 4C, upper panels), this effect appeared to be obscured in right abducens nerves of one-eared animals (Figure 4B and 4C, lower panels). Discharge modulation of these nerves were found to exhibit a considerable heterogeneity, with some nerves modulating with profiles that were entirely inconsistent with typical right abducens nerves of control animals. Quantification of phase relationships with respect to leftward head motion velocity (Figure 4D-E) was in agreement with such qualitative observations. Indeed, abducens nerves in controls were found to exhibit temporal-activity patterns consistent with those expected for their anatomical identity (Figure 4D), with no temporal overlap of the activity in their bilateral abducens counterparts (see blue hashed bars in Figure 4D). Directional phase analysis *re* leftward velocity revealed mean phase vectors of 143.92° ±13.52° (r = 0.973) and 329.86° ±23.48° (r = 0.919) for left and right abducens, respectively. These activity metrics are demonstrative of preferred directional firing, which indicates spatially separate tuning properties present in these nerves (Figure 4E; *p* < 0.0001 and *p* = 0.000067, Rayleigh’s Uniformity Test for left and right abducens, respectively; *p* < 0.001, Moore’s Paired Test). In contrast, left and right abducens nerves in one-eared animals showed a large spread in temporal distribution of activity during rotation (Figure 4C-D), with mean directional vectors for the left and right abducens nerves of 152.77° ±21.24° (*r* = 0.934) and 282.16° ±102.02° (*r* = 0.205), respectively (Figure 4E). In particular, right abducens nerves appeared to modulate in some cases even during rightward peak velocity (Figure 4C-D) and failed to exhibit a preferred directional sensitivity (*p* = 0.614, Rayleigh’s Uniformity Test). Conversely, left abducens nerves from one-eared animals largely exhibited an appropriate temporal pattern of discharge modulation and grouped in a preferred direction (*p* < 0.001, Rayleigh’s Uniformity Test). These results suggest a clear lack of entirely separate spatial tuning between both nerves (*p* > 0.05, Moore’s Paired Test). The mean angular directional preferences for left abducens nerves between controls and one-eared animals were found to be no different (*p* = 0.365, Watson-Williams F-test), as well as for comparison of right abducens nerves (*p* = 0.241, Watson-Williams F-test), suggesting that in some animals appropriate tuning properties are present, despite the heterogeneity introduced by individual animals.

Amplitude-dependent features, such as the depth of modulation, were found to be not statistically different between anatomical left and right nerves within controls and manipulated animals (Supplemental figure 3A_1_-A_2_; *p* = 0.1563, *p* > 0.9999; Wilcoxon signed-rank test, *n* = 6 pairs of controls and *n* = 5 pairs of one-eared animals, respectively). Such features are characteristic of a spatially appropriate push-pull aVOR organization (Straka and Dieringer, 2004). In addition, the spontaneous activity, corresponding to discharge rates during periods of no head motion, were similarly invariant between left and right nerves in both animal groups (Supplemental figure 3B_1_-B_2_; *p* = 0.5625, *p* = 0.6250; Wilcoxon signed-rank test, *n* = 6 pairs of controls and *n* = 5 pairs of one-eared animals, respectively), suggesting the presence of a homeostatic plasticity during the ontogenetic establishment of the circuitry that apparently aims at symmetric driving forces. Comparison of discharge rates during the application of rotational stimuli relative to spontaneous resting activity rates (modulation index) revealed expected response profiles in control nerves (Supplemental figure 3C). Left and right abducens nerves appeared to modulate around their spontaneous firing rate, with frequencies modulating below and above their resting rate, corresponding to stimulus-evoked periods of discharge facilitation and disfacilitation (Supplemental figure 3C, blue heat maps). Modulation around resting activity was observed less often in one-eared animals, in both left and right nerves, suggesting that at least in some cases these nerves in one-eared animals suffer from a lack of appropriate facilitation/disfacilitation dynamics based on the activity pattern from the singular ear (Supplemental figure 3C, orange heat maps). Between control and one-eared animals, the modulation depths in both nerves were significantly less robust (Supplemental figure 3D_1,_ D_2_; *p* = 0.0463, *p* = 0.0011; Mann-Whitney *U*-test for left and right abducens nerves, respectively), while resting rates were found to be no different (Supplemental figure 3D_1,_ D_2_; *p* = 0.3823, *p* = 0.1876; Mann-Whitney *U*-test for left and right abducens, respectively). Beyond differences in modulation depth, these physiological profiles suggest a striking dissimilarity in bilateral abducens nerve activity during the aVOR for animals which have developed with only a singular ear. Motor transformations from such singular ears appear to follow temporal dynamics of comparable extent as would be expected in control unmanipulated animals for anatomically defined left abducens nerves. This is not too surprising given the major driving force of abducens motor nerve activity from the contralateral ear, which is the residual singular ear in one-eared animals. In contrast, right abducens nerves exhibit a prominent temporal heterogeneity. The disparity between the two nerves, which in canonical conditions does not exist, is indicative of potentially equally heterogenic mechanisms used to permit modulatory activity that is necessary to yoke the eyes.

## Discussion

Unilateral extirpation of the embryonic otic placode generated tadpoles that developed with a singular ear. These one-eared tadpoles exhibited a considerable degree of developmental plasticity, observable during execution of the horizontal vestibulo-ocular reflex. Eye movements, though weaker compared to two-eared controls, demonstrated successful execution of sensorimotor transformations despite the lack of bilateral mirror-symmetric vestibular endorgans. Achievement of this capacity occurs through neuronal computations of inputs from the singular inner ear in hindbrain vestibular centers. Once in the hindbrain, input from the single ear is sufficient to drive bilateral directed gaze stabilizing reflexes. The developing central nervous system is therefore capable of establishing directionally sensitive sensorimotor processing capabilities from self-motion information originating from a single set of vestibular endorgans. The mechanisms driving this capacity likely derive from individualized strategies of circuit plasticity during development which are largely independent of visual-motion contributions.

### Developmental plasticity in unilateral sensory deprived vestibulo-ocular circuits

Surgical excision of the otic placode at early stages in *Xenopus laevis* generated embryos which experienced the complete absence of one inner ear and were thus challenged with detecting self-motion stimuli with only one set of endorgans. Downstream of such challenges in sensory detection, integration of vectorial inputs through non-canonical peripheral pathways was continued, despite the fact that these pathways typically receive bilateral motion vectors (Glasauer and Knorr, 2020). Unilateral ablation techniques such as this have previously been demonstrated as a suitable approach to assess the effects of sensory deprivation on hindbrain targets in *Xenopus* (Fritzsch, 1990; Elliott et al., 2015a, 2015b), chick (Levi-Montalcini, 1949; Peusner and Morest, 1977), and salamanders (Goodman and Model, 1988). While such studies were pivotal in identifying the effects on central vestibular circuit development, the extent and effective execution of sensorimotor transformations from the remaining singular inner ear by vestibular-ocular motor centers is so far unexplored. Related embryonic manipulations such as surgical rotation (Lilian et al., 2019; Elliott et al., 2015b) or addition of supernumerary ears (Elliott et al., 2015a; Gordy et al., 2018), profiled functional achievements to a successful degree, though central computations retain inputs of variable degrees and spatio-temporal composition from both sides (Lilian et al., 2019; Elliott et al., 2015a,b).

Beyond different surgical techniques, non-invasive methods have been used to profile vestibular sensory loss, particularly from selective deficiencies in microgravity (Horn, 2003), inner ear genetic manipulations with permanent effects (Kopecky et al., 2012; Macova et al., 2019) or those of a more transitory nature, such as the generation of Zebrafish with temporary utricular deprivation (Roberts et al., 2017; Ehrlich and Schoppik, 2019). Such manipulations, however, were either not specific to one side (Kopecky et al., 2012), lack uniformity of deprivation across all sensory epithelia (Roberts et al., 2017), or present with defects in various sensorimotor areas (Patten et al., 2012). Functional consequences of these conditions would therefore derive from motion information of both sides, albeit with varying degrees of asymmetric signaling. In contrast, one-eared animals in the current study receive self-motion information solely through one inner ear, which lacks its bilateral mirror-symmetric compliment, but normally develops all other sensorimotor systems, with minimal detrimental effects on adjacent placode-derived sensory organs (Elliott et al., 2010). The overall retention of vestibulo-motor responses in the presence of a singular ear (Figure 1) demonstrates the capacity of one-eared animals to execute adequate spatio-temporal VOR transformations. Execution of gaze stabilizing vestibular reflexes in darkness, and thus without visually derived motion-signaling (Straka and Dieringer, 2004), demonstrate that these animals have generated sufficient plastic vestibular alterations to transform directionally specific inputs from the singular ear. Such plastic capabilities have not been observed in previous behavioral assessments of one-eared *Xenopus* tadpoles (Zarei et al., 2017), where Mauthner-cell mediated swimming startle responses were of appropriate measure, although directionally biased with respect to inputs from the singular ear (Zarei et al., 2017). The latter finding is not entirely surprising, given the physiological basis of Mauthner cell-mediated startle behaviors (Korn and Faber, 2005), where no morpho-physiological modifications within the singular ear can obviously encode bidirectional stimuli. In contrast, in the current study, lateralized horizontal rotation is detectable by the singular horizontal semicircular canal and was shown to derive from the structurally guided facilitation/disfacilitation dynamics of semicircular canal afferent signals (Figure 2).

Particularly surprising was the unexpected inequality in eye motion amplitudes during contraversive *versus* ipsiversive (with respect to the single ear) rotation, which favored more robust responses during disfacilitation of the singular right ear. Acute lesion of a single stato-acoustic nerve in *Xenopus* tadpoles showed a physiologically more expected effect where rotation toward the lesion side elicited very poor eye movements, a feature consistent with the sudden loss of a predominant directional sensitivity (Soupiadou et al., 2020), which is likely due to the resulting absence of the driving force supplying relevant extraocular motoneurons (Branoner and Straka, 2018). Here, despite the obvious bidirectional sensitivity of the singular inner ear, such asymmetric motor output highlights individualized differential strengths in computation within brainstem processing regions. This suggests that plasticity mechanisms are not goal-directed at consistently aiming for production of suitable motor commands that equalize sensitivity vectors (Dieringer, 2003). However, behavioral responses during oscillation-driven facilitation and disfacilitation of singular ears seems to provide sufficient dynamics for the production of spatio-temporally appropriate aVOR responses beyond differences in directional vectors. In animals with an acute loss of inputs from one inner ear, the residual oscillatory motion-driven aVOR is much less robust and generally rather asymmetric with a predominance of responses during rotations toward the intact side (Soupiadou et al., 2020).

In the visual system, early reversible monocular deprivation in the cat leads to an increased responsiveness to signals from the remaining eye in cortical areas, with a concomitant expense of target sensitivity to inputs from the shunted eye (Wiesel and Hubel, 1963). The data presented here suggest the opposite, with a dampening in motion vector sensitivity in the canonical excitatory direction of the singular ear. However, the marginal redundancy in visual input originating from individual eyes during visual motion detection, even among lateral- and frontal-eyed animals which present with markedly strong directional asymmetries (Masseck and Hoffmann, 2009; Wagner et al., 2022) does not exist for vestibular signal encoding and processing, where mirror-symmetric endorgans encode directional domains that are almost mutually exclusive. Therefore, that the remaining inner ear maintains and potentially increases the ability to peripherally distinguish directional vectors (Figure 2) suggests that one-eared tadpoles have individually activated developmental strategies that ultimately provide preservation or extension of bidirectional sensitives. Two-eared mediated bilateral modulation of motion-related neuronal activity is known to depend on spontaneous afferent discharge levels, which provide a larger range for bidirectional motion encoding at higher resting rates and a more directionally restricted sensitivity at low or very low afferent firing rates as usually present in amphibian species (Blanks and Precht, 1976; Honrubia et al., 1981, 1989; Straka, 2020). Here, a generalized strategy to generate bidirectional sensitivity from singular inner ears might involve the establishment of higher resting discharge rates in vestibular afferent fibers beyond the usually low firing rates to allow encoding of head motion only in the on-but not in the off-direction. Developmentally established higher resting rates of vestibular afferents innervating the singular ear would thus considerably extend the dynamic range for the motion encoding by increasing the degree for a firing rate disfacilitation during contraversive head movements.

One-eared *Xenopus* tadpoles subjected to drop-swim assays showed deficits in postural stabilization (Elliott et al., 2015b), which suggests an inability to correct for directionally asymmetric vestibular inputs. However, the extent to which this reflects a limitation in processing bandwidth required for integrating otolith and semicircular canal inputs, or is merely a developmental restriction, given that tadpoles were assessed relatively shortly after the ear removal, remained untested (Elliott et al., 2015b). Developmental progression of similarly manipulated tadpoles to the physiological stages assayed here has been done previously but was limited to tract tracing observations alone (Fritzsch et al., 1990). The capability of one-eared animals in this study to execute a spatially appropriate aVOR provides a unique perspective on the developmental strategies for adaptive plasticity, highlighting the extent to which directional sensitivities may develop despite lacking canonical structures for their detection. These results expand upon the observed persistency of appropriate vestibular processing despite embryonic deficiencies in peripheral inputs. Indeed, delayed bilateral otolith formation in Zebrafish demonstrated a similar autonomy for nascent posture-stabilizing circuits (Roberts et al., 2017). The extent of developmental plasticity observed in the current study compliments with previous experimental models in the visual system of amphibians (e.g., Constantine-Paton and Law, 1978; Ruthazer et al., 2003; Blackiston et al., 2017) and teleosts (e.g., Ramdya and Engert, 2008), which served to highlight the remarkable degree of flexibility during development of sensory systems. A potential mechanism in the case of one-eared tadpoles might include altered resting firing rates as well as a shift in the push-pull organization of inhibitory and excitatory vestibulo-ocular connections. Such mechanisms could generate a spectrum of individually specific encoding capacities for bilateral extraocular motor commands through alterations in the degree of excitation or disinhibition (see below).

### Mechanisms of developmental vestibular plasticity

Firing activity of extraocular motor nerves represent the terminal site of VOR sensorimotor transformations originating from inner ear peripheral inputs (Gensberger et al., 2016). Extracellular discharge dynamics of these motoneurons, particularly those of the abducens nerve, have previously been used to profile downstream circuit computations after an acute vestibular loss in *Xenopus* (Lambert et al., 2013; Branoner and Straka, 2018), and various species of *ranid* frogs (e.g., Rohregger and Dieringer, 2003), as well as following embryonically guided introduction of additional vestibular inputs (Gordy et al., 2018). Here, profiling abducens nerve dynamics in one-eared animals reported on a considerable spectrum of developmental plasticity measures. Despite the absence of one inner ear, a sustained and robust spontaneous resting rate of the extraocular motor nerve was observed. The presence of such robust rates contrasts with animals following an acute unilateral vestibular loss where an elimination of resting activity in extraocular motor nuclei contralateral to the lesioned ear was reliably demonstrated (Branoner and Straka, 2018; Lambert et al., 2013). Given that motoneurons of the right abducens nerve in one-eared tadpoles exhibit such prominent resting rates despite lacking a contralateral left inner ear, which under control conditions provides the excitatory drive (Straka and Dieringer, 1993), suggests the presence of homeostatic mechanisms which likely aim at establishing symmetric driving forces during ontogeny. Following an acute lesion of one stato-acoustic nerve at the tadpole stage (e.g., Lambert et al., 2013), such a loss is likely driven and permanently maintained by the weighted inputs on abducens motor targets from second-order vestibular neurons which suddenly lack excitatory inputs from the lesioned side while maintaining continued ipsilateral inhibition from the remaining ear. In the current study, the development of a suitable driving force could likely be generated by both a decrease in inhibitory inputs to the right abducens nucleus as well as by an indirect excitatory input from the remaining inner ear. Unilateral labyrinth-ectomized *ranid* frogs appear to rely heavily on the former compensatory strategy, though were rather heterogeneous in the efficacy of their responses (Agosti et al., 1986).

In the current one-eared animals, the impaired ability of right abducens nerves to modulate around their respective resting rates in some animals suggests a degree of inadequacy in disinhibition and might lend support to this notion. Indirect excitatory contributions might be the result of midline crossing commissural pathways in the hindbrain (Straka, 2020), particularly of excitatory fibers which innervate horizontal semicircular canal second-order vestibular neurons (Holler and Straka, 2001) and assist the generation of symmetric resting rates, as might be the case for the acute loss in *ranid* frogs (Agosti et al., 1986). The relative synaptic weights of such excitatory connections, their second-order targets, and the distributions relative to inhibitory commissural fibers is thus of great interest to investigate. The behavioral delay in aVOR-related eye movements of the current study tends to support such a claim, given the long latency to reach peak eye motion velocity relative to control conditions and could be due to additional synaptic relays during sensorimotor transformation (Figures 1, 2). Commissural pathway-mediated generation of such symmetry would be opposite to that observed in cats, where a loss of crossed vestibular commissural inhibition causes an increase in the resting discharge of contralesional second-order vestibular neurons, which, however, decreases over time (Yagi and Markham, 1984). In chicks, an increase in excitatory inputs on the lesioned side was found only in animals which had not been classified as being able to behaviorally compensate for an acute unilateral vestibular loss (Shao et al., 2012). The similar resting rates between bilateral abducens nerves in embryonically manipulated *Xenopus* tadpoles approximate a considerable extent of symmetric activity in their upstream vestibular nuclei. These animals, though lacking behavioral and physiological robustness at control levels, have seemingly developed such a symmetry, which permits appropriate motor output and highlights the general need of symmetric activity levels in vestibular nuclei, as has been proposed in several experimental models (Lambert and Straka, 2012).

Motion evoked discharge rates in the abducens nerves of one-eared tadpoles were cyclic with respect to the stimulus. Despite differences in the ability to modulate around their respective resting rates, abducens activity profiles clearly demonstrated a general capability to execute sensorimotor transformations originating from inputs from the singular inner ear. However, the notable heterogeneity in the response phase of individual nerves indicates a range of temporal relationships. This is particularly evident for right abducens nerves, where peak firing rates temporally extended even in some cases to periods with inappropriate motion direction. In these abducens motoneuron populations, the lack of direct excitatory input from the operated side, despite disinhibitory contributions from the remaining ear, might seem to be a detriment that was sometimes developmentally uncompensated for, particularly given that all left abducens nerve responses appeared appropriate in phase (with excitatory inputs from the residual singular inner ear). However, the behavioral data suggests against this, particularly given the activation of considerably strong eye movements during sinusoidal and unilateral motion toward the operated side. Therefore, right abducens motoneurons with directionally inappropriate phase metrics might be supplemented with temporally complimenting, though canonically phase shifted, discharge rates in medial rectus-innervating oculomotor motoneurons, which would provide suitable antagonistic yoking required for the aVOR (Horn and Straka, 2021). Post-lesional plasticity in *ranid* frogs has so far demonstrated a considerable variability in the spatial tuning of the abducens nerve activity during linear and angular motion-evoked VOR, which was demonstrated to be behaviorally detrimental but likely beneficial for the survival of the deafferented central neurons (Goto et al., 2001; Rohregger and Dieringer, 2003; Dieringer, 2003). In tadpoles of the current study, the ability to execute spatially meaningful aVOR behaviors suggests that inappropriate tuning of abducens nerve activity might only play a minor role. Precise tuning characteristics of central vestibular neurons would be beneficial to further explore such relationships, such as in recent approaches quantifying tuning and convergence properties in Zebrafish (Liu et al., 2020)

Motion-sensitive sensory modality integration is prominent in brainstem gaze and posture processing centers (Angelaki and Cullen, 2008), and plasticity-based reorganization following vestibular loss is typically supplemented by these modalities (Curthoys, 2000). In tadpoles of the current study, concurrent optokinetic flow appeared to not supplement aVOR responses neither in amplitude nor in temporal attributes. These animals have therefore developed a vestibular processing regime without relying on augmented synergistic visual motion signals, which suggests the location of plasticity as being possibly exclusive to vestibular circuit elements alone. A wealth of studies has reached similar or contrasting conclusions, which highlights broad species differences in the apparent extent of modality substitution following vestibular deprivation (Dieringer, 1995; Darlington and Smith, 2000). The current study is thus the first instance of embryonic vestibular deprivation demonstrating the impact on the performance of vestibulo-ocular reflexes that align with independence from visually mediated substitution. Cerebellar contributions to developmental maturation of vestibular evoked posture-stabilization (Ehrlich and Schoppik, 2019), as well as homeostatic mechanisms following prolonged rotation (Dietrich and Straka, 2016) implicate the possibility of the cerebellum as being involved in plasticity strategies here as well, though experimental validation is still pending. Ontogenetic development of brainstem vestibular circuits is therefore highly plastic and can be exploited to drive functionally appropriate motor outputs despite lacking canonical peripheral sensors. Considerations to such plasticity extents would be beneficial for targeted therapeutics, such as those aimed at using transplantation approaches to replace vestibular deficits (Elliott et al., 2022), and might aid in the holistic understanding of adaptability in vestibular development and processing.

## Acknowledgements

The authors thank Michael Forsthofer for his insightful and constructive comments on earlier versions of this manuscript as well as Dr. François Lambert, Gabriel Barrios, Michael Forsthofer, and Prof. Dr.-Ing. Stefan Glasauer for guidance with the analysis and scripts. Gratitude is also due to the LMU Biocenter Animal Facility veterinarian staff for their assistance in rearing experimental animals. Confocal microscopy was performed in the “Center for Advanced Light Microscopy” (CALM) facilities of the LMU Munich. This research was supported by the Deutsche Forschungsgemeinschaft (German Science Foundation; CRC 870, RTG 2175, STR 478/3-1).

## Author Contributions

Conceptualization, C.G. and H.S.; Methodology, C.G. and H.S.; Software, C.G.; Validation, C.G.; Formal Analysis, C.G.; Investigation, C.G.; Resources, H.S.; Data Curation, C.G and H.S.; Writing – Original Draft, C.G. and H.S.; Writing – Review and Editing, C.G. and H.S.; Visualization, C.G.; Supervision, H.S.; Project Administration, H.S.; Funding Acquisition, H.S.

## Methods

### Animals

Experiments were conducted on *Xenopus laevis* embryos and larvae at different developmental stages (Nieuwkoop and Faber, 1994) and either sex. Embryos were obtained through induced ovulation by injection of human chorionic gonadotropin, followed by *in vitro* fertilization with sperm suspension in 1 x Modified Barth’s Saline (MBS, diluted from 10 x stock; 880 mM NaCl, 10 mM KCl, 100 mM HEPES, 25 mM NaHCO_3_, pH 7.6) or manual collection after natural mating. Embryos from either fertilization method were de-jellied with 2% cysteine and incubated in 0.1 x Marc’s Modified Ringer’s Solution (MMR, diluted from 10 x stock; 1 M NaCl, 18 mM KCl, 20 mM CaCl_2_, 10 mM MgCl_2_, 150 mM HEPES, pH 7.6-7.8) until animals reached stage 46, when tadpoles were transferred into standing tanks of de-chlorinated water of appropriate volume (McNamara et al., 2018), maintained at 17-19°C under a 12 hour/12 hour light/dark cycle, and fed daily with a powdered Spirulina (Algova, Germany) suspension in tank water. After reaching stage 53-57, tadpoles were used for behavioral, anatomical and/or physiological assessment in accordance with the “Principles of animal care” publication No. 86–23, revised 1985, of the National Institutes of Health and were carried out in accordance with the ARRIVE guidelines and regulations. Permission for the experiments was granted by the legally responsible governmental body of Upper Bavaria (Regierung von Oberbayern) under the license codes ROB-55.2-1-54-2532-14-2016, ROB-55.2.2532.Vet_03-17-24 and ROB-55.2.2532.Vet_02-19-146. In addition, all experiments were performed in accordance with the relevant guidelines and regulations of the Ludwig-Maximilians-University Munich.

### Inner Ear Extirpation

Extirpations of the inner ear anlage (the otic placode) were performed in 1.0 x MMR at a room temperature of 22°C on stage 25-27 embryos. Embryos were anesthetized with 0.02% Benzocaine (Elliott and Fritzsch, 2010) prior to the surgical manipulations. All surgical interventions were performed with fine tungsten needles (0.125 mm, Fine Science Tools, 10130-05). Access to the developing inner ear following visual identification of the target area was made by peeling back the dorsolateral ectoderm-derived layer overlying the developing otic placode. Placodes were subsequently identified by visual inspection and were surgically excised from the surrounding tissue (Supplemental video 1). Removals were done unilaterally, with the contralateral side left unmanipulated. Care was taken to minimize the ablation and disturbance of adjacent non-otic tissue such as to exclusively remove the developing ear. After the surgery, embryos were maintained for 30 minutes in 1.0 x MMR to permit healing of the exposed tissue before being returned to 0.1 x MMR and reared until reaching the desired stages for the different types of experiments (see below). A representative video of ear extirpation was captured on a SteREO Discovery.V20 stereo microscope with a Axiocam 305 color camera (Carl Zeiss Microscopy GmbH) taken at 8 fps and exported at 15 fps using ZEN software 3.4.91 (Carl Zeiss Microscopy GmbH).

### Experimental preparations

All experiments were performed on semi-intact *in vitro* preparations generated from tadpoles that had been subjected to a unilateral embryonic inner ear extirpation or from untreated control animals and were obtained following a protocol described previously (Soupiadou et al., 2020; Knorr et al., 2021). Tadpoles were first anesthetized in 0.05% 3-aminobenzoic acid ethyl ester methanesulfonate (MS-222; Pharmaq Ltd. UK) at a room temperature of 22°C for 3-5 min and were then transferred into ice cold frog Ringer solution (75 mM NaCl, 25 mM NaHCO_3_, 2 mM CaCl_2_, 2 mM KCl, 0.1 mM MgCl_2_, and 11 mM glucose, pH 7.4). An *in vitro* preparation was generated by decapitation, removal of the lower jaw, and evisceration. The skin covering the dorsal part of the head including the otic capsule(s) was removed, the cartilaginous skull opened and the choroid plexus detached to allow access of the Ringer solution to the brainstem through the open fourth ventricle. Such *in vitro* preparations maintain fully functioning sensory organs (e.g., ears and eyes) as well as all central nervous circuits, and contain intact peripheral motor nerves and effector organs (e.g., extraocular muscles; see Straka and Simmers, 2012). Following surgical procedures, animals were allowed to recover for 2 hours at 17°C.

### Visuo-vestibular motion stimulation

Vestibular sensory stimulation was provided by a six-axis motion stimulator (PI H-840, Physik Instrumente, Karlsruhe, Germany) mounted onto a breadboard table (TMC Ametek). Semi-intact tadpole preparations were mechanically secured with insect pins in the center of a Sylgard-lined chamber (Ø 5 cm) and continuously superfused with oxygenated (Carbogen: 95% O_2_, 5% CO_2_) Ringer solution to maintain a constant temperature of 17.5 ± 1.0°C. For behavioral experiments, horizontal sinusoidal motion stimuli were generated by a custom written software in C++ (Soupiadou et al., 2020) and delivered to the control unit of the motion stimulator. Stimulation paradigms consisted for each animal as follows: bouts of sinusoidal vestibular stimulation were provided through oscillating horizontal rotation performed at 0.5 Hz with a peak velocity of ±31.4°/s for 15 consecutive cycles, followed by an inter-stimulus period of at least 60 seconds. Each animal was provided with the horizontal rotational stimuli first in darkness and then in light, with the light condition corresponding to motion in the presence of a world-stationary visual scene consisting of black and white stripes used for optokinetic stimulation described below. In both darkness and light, stimulation bouts were initiated with motion beginning either in the leftward or rightward directions prior to oscillation at 0.5 Hz (2 second period) between both directions, with leftward initiating bouts occurring first in the order of presented stimulus paradigms before those starting rightward. In all bouts, the first half cycle (1.0 second) of each bout were classified as unidirectional stimulation for subsequent analyses. Optokinetic stimuli were generated by three digital light processing video projectors (Aiptek V60). Visual patterns were projected onto a cylindrical screen (Ø 8 cm, height 5 cm) positioned around the center of the motion platform, providing a 275° visual field with a refresh rate of 60 Hz. Patterns consisted of equally spaced vertically oriented black and white stripes of 16°/16° spatial size. Optokinetic stimuli were presented at three frequencies, initiating in the following order: 0.1, 0.2 and 0.5 Hz, and occurred in 3 repetitions of 15 consecutive cycles per frequency, interrupted by a stationary period of at least 15 seconds. Vestibular stimulation paradigms for electrophysiological recordings of extraocular motor nerve discharge were performed similarly with consistent sinusoidal parameters, however stimulation was performed only in darkness, and consisted of at least two stimulation bouts of 5-15 cycles each (at 0.5 Hz with a peak velocity of ±31.4°/s as indicated above). Vestibular and optokinetic stimulus profiles were set to be sampled into Spike2 signal recording software (Cambridge Electronic Design, UK) at a rate of 50 Hz.

### Eye Motion Tracking

Eye movements, in response to head motion (vestibular) and exclusive visual image motion (optokinetic) stimulation, were recorded by a digital camera (Grasshopper Mono, Point Grey Research Inc., Canada) fitted with a high-pass infrared filter lens and appropriate zoom objective (Optem Zoom 70XL, Qioptiq Photonics GmbH & Co. KG, Germany; M25 × 0.75 + 0.25). The camera was mounted onto the motion simulator platform and centered directly above the semi-intact preparation. Video sequences were recorded with a frame capture rate of 30 Hz using the FlyCap2 software (v2.3.2.14.) under illumination of the *in vitro* preparation with an infrared light source. Positional changes of both eyes over time were quantitatively assessed (Beck et al., 2004) by fitting an ellipse to each eye independently and computing the deviation of the major axis of each ellipse from the longitudinal image axis in each video frame. Behavioral data were captured and digitized at 30 Hz by a CED 1401 A/D interface and associated Spike2 program (Cambridge Electronic Design Ltd., United Kingdom).

### Electrophysiological recordings of extraocular motor nerves

Extracellular multi-unit spike discharge was recorded from the severed ends of the *lateral rectus* (LR) motor nerves close to the innervation site of their respective bilateral eye muscles with glass suction electrodes. Electrodes were made from glass capillaries (Science Products, GB100-10) with a horizontal puller (P-87, Sutter Instruments Co., USA) and were individually broken to fit to the size of each nerve. Multi-unit spike activity was recorded (EXT 10-2F; npi electronic GmbH, Germany) and digitized at 20 kHz by the CED 1401 A/D interface in Spike2.

### Immunohistochemistry

Young tadpoles at stage 46 were anesthetized in 0.5% MS-222 and fixed by immersion in 4% paraformaldehyde (PFA) in phosphate-buffered saline (PBS) for at least 3 hours at 4°C. Following, tadpoles were dissected by removal of the lower jaws and viscera, decapitated at the head/tail junction, and freed from the skin overlying the dorsal head. Subsequently, samples were dehydrated in 70% ethanol for a period of 1 - 12 hours, followed by washing 3 x in 0.1x PBS before immersion for 1 hour at a room temperature of 22°C in 5% normal goat serum with 0.1% Triton X100 to block the immunoreactive epitopes. Samples were then incubated overnight at 36°C with primary antibodies against the hair cell marker Myosin VI (1:400; Proteus Biosciences, 25-6791) and the neuronal marker acetylated-tubulin (1:800; Sigma-Aldrich, T7451). Afterwards, washing and blocking reactions were repeated as above, prior to incubation for 1 hour at room temperature with species-specific secondary antibodies (1:500, Alexa) and DAPI (1:1000) for 1 hour at room temperature. Following a series of washes (6x) in 0.1x PBS, animals were mounted on microscope slides, coverslipped with Aqua Polymount (Polyscience) and subsequently imaged on a Leica SP5-2 confocal microscope.

### Data Analysis

Data analysis of eye motion and extraocular motoneuron spike discharge recordings were conducted post-hoc in Python 3 following export from the Spike2 acquisition software into MATLAB (The Mathworks, Inc.) data files. Eye positions for both eyes were re-sampled at 200 Hz and low-pass filtered with a cut-off frequency of 4 Hz (Butterworth; 2^nd^ order) and then averaged together. Individual sinusoidal stimulus cycles, which were determined to contain episodes of stimulus-unrelated eye twitches, corresponding to fast-phases or other spontaneously occurring eye motions, were manually identified and removed from subsequent analysis (Beck et al., 2004). Responses evoked by individual sine waves were averaged across multiple cycles within each animal. The general lack of visuo-vestibular motion stimulus-driven resetting fast phases in *Xenopus* tadpoles at the tested stimulus settings facilitated the calculation of response gains as: the ratio of peak-to-peak eye position to peak-to-peak stimulus position, and corresponding phase metrics as: temporal delay between peak stimulus position and peak eye position calculated as an angular fraction of the motion cycle. Peak amplitudes during unidirectional stimulation were calculated during the first half cycle (1 second) initiating a stimulation bout, corresponding to the peak eye motion response across this period.

Extraocular motor discharge data was filtered with a Butterworth bandpass filter with lower and upper limits of 200 and 600 Hz, respectively, to reduce noise generated by the platform motion. Discharge rates were calculated from spike counts over time following a manual amplitude and spike interval-dependent threshold selection, which was determined for each individual nerve in each animal. Spike counts during each stimulus cycle were used to produce a peri-stimulus time histogram (PSTH; bin size 0.05 seconds) over a single cycle. For PSTH generation, stimulus cycles were selected from sinusoidal bouts with respect to peak directional velocity contralateral to each nerve to better identify the temporal dynamics. Spike rates were then calculated by first dividing spike counts within each histogram bin by the number of cycles and then by bin size. Responses which contained episodes of stimulus-unrelated eye twitches were excluded manually. For visualization of PSTHs in heat maps, responses were either normalized to their respective peak discharge rate per individual animal or as raw rates as indicated. Resting nerve discharge rates were calculated from the average of 20 seconds of spontaneous activity during periods where the head remained stationery either prior to and/or between stimulus bouts. Discharge rates over a single motion cycle were used to calculate parameters of modulation depth and phase, corresponding respectively to the difference between the highest and lowest rate in the former, and the angular fraction of the difference between peak discharge rate and peak stimulus directional velocity in the latter. This latter calculation was also done with respect to opposite velocity motion for each nerve by shifting the angular location of peak firing rate forward by 180° along a 360° scale prior to determining the angular difference as above. Owing to unequal sampling rates in stimulus motion position, stimulus velocity metrics were acquired by re-sampling a single positional cycle to 20 kHz and fitting to a sine wave prior to differentiation.

### Statistics

Statistical differences between independent (two-eared control *versus* one-eared animals) data sets were assessed using the Mann-Whitney *U*-test for unpaired nonparametric data, and the Wilcoxon matched-pairs signed-rank test for paired (within experimental groups) nonparametric data in Prism (GraphPad Software 8.4.3, Inc, USA). Gain and phase comparisons for tadpoles across multiple frequencies were performed with the nonparametric Friedman test followed by a Dunn’s multiple comparisons test. Circular statistics for electrophysiological data was calculated in Oriana (Version 4.02; Kovach Computing Services) as shown previously (Bacqué-Cazenave et al., 2018). Pooled phase values *re* peak leftward velocity (see above) taken from left and right abducens nerves from individual animals were used to calculate a mean vector, defined by an angular direction in degrees (*μ*, ± circular standard deviation) and a corresponding length metric approximating clustering strength around the mean (*r*). Assessment of uniform distribution, indicative of no preferred direction, was calculated by Rayleigh’s Uniformity Test (*p*). Significance of difference between mean angular directions from pooled left and right abducens nerves in each animal group was tested with Moore’s Paired Test. Differences between mean angular directions in control and one-eared animals was assessed with pairwise Watson-Williams F-test. A significance threshold of 0.05 was used for all analyses. Population data is reported as mean ± standard deviation (SD) unless otherwise stated.

## SUPPLEMENTAL MATERIAL FIGURE LEGENDS

**Supplemental video 1. Embryonic extirpation of the otic placode.** Representative procedure of a unilateral removal of the inner ear anlage in a stage 26 *Xenopus laevis* embryo.

**Supplemental figure 1. Phenotype validation of one-eared tadpoles. (A-D)** Representative unmanipulated control **(A_1_**-**B_3_)** and left-side otic placode-extirpated **(C_1_**-**D_3_)** stage 46 tadpoles, specifically depicting the two bilateral and the one unilateral inner ear(s), respectively; whole-mount immunohistochemical labeling of control **(B_1_**-**B_3_)** and one-eared **(D_1_-D_3_)** tadpoles stained for the nuclear marker DAPI (gray; **B_1_**, **D_1_**) revealed the structure of the inner ears **(B_1_, D_1_** right) and the cranial otic region in the absence of an ear (white * in **D_1_)**; neuronal marker, acetylated tubulin (green), and hair cell/muscle marker, myosin-VI (red), identifies **(B_2_**-**B_3_**, **D_2_-D_3_**) hair cell clusters and corresponding VIII^th^ nerve innervations, indicative of inner ear endorgans **(B_3_)**, which are entirely absent from the corresponding region on the left side of one-eared animals **(D_3_)**; **B_1_** and **D_1_** are single color channels of **B_2_**, **D_2_**, respectively; white boxes in **B_2_**, **D_2_** approximate the region of higher magnification shown in **B_3_**, **D_3_**. **(E**, **F)** Representative stage 55 control **(E_1_-E_2_)** and one-eared **(F_1_-F_2_)** animals depicting the absence of an otic capsule and inner ear endorgans on the left side; **E_2_** and **F_2_** are higher magnifications of images in **E_1_**, **F_1_**; Scale bars are 1 mm in **A_1_**-**A_2_** and **C_1_-C_2_**, 100 µm in **B_1_**-**B_3_** and **D_1_**-**D_3_** and 2 mm in **E_1_**-**E_2_** and **F_1_**-**F_3_**. R, rostral; C, caudal; M, medial; ac, hc, pc, anterior, posterior, horizontal semicircular canal cristae; ut, sc, utricular, saccular macula; bp, basilar papilla; aLL, pLL, anterior, posterior lateral line nerve; III, oculomotor nerve; V, trigeminal nerve; VII, ‘facia’ nerve; VIII; stato-acoustic nerve; IX, glossopharyngeal nerve; X, vagus nerve; Tel, telencephalon; OT, optic tectum; Hb, hindbrain; SC, spinal cord.

**Supplemental figure 2. Optokinetic reflex performance in one-eared tadpoles. (A, C)** Schemes of the experimental approach depicting sinusoidal horizontal rotation of a patterned black- and white-striped visual scene (±10° movement excursion) while the head is maintained stationary in controls (**A**) and one-eared animals (**C**). **(B, D)** Averaged responses over a single cycle of visual image motion at 0.1 Hz (left, peak velocity: 6.28°/s), 0.2 Hz (middle, peak velocity: 12.57°/s) and 0.5 Hz (right, peak velocity: 31.4°/s) for controls (**B**; *n* = 13, calculated from 21-45 cycles) and one-eared animals (**D**; *n* = 12, calculated from 26-45 cycles). Blue- and orange-colored lines and shading indicate population means ±SD. Solid black lines indicate the visual scene motion stimulus; individually calculated gain **(E-F)** and phase *re* visual stimulus position **(G-H)** for all three stimulus frequencies of each experimental group (**E, G**; left, control; right, one-eared animals) and comparison between experimental groups. Significance levels are indicated by asterisks: * *p* ≤ 0.05, *** *p* ≤ 0.001, **** *p* ≤ 0.0001 (Friedman nonparametric test for matched pairs with Dunn’s multiple comparison test in **E, G**, Mann-Whitney *U*-test in **F, H**). Horizontal dotted lines in **G-H** at 0° indicate phase alignment of the stimulus.

**Supplemental figure 3. Multi-unit discharge dynamics in abducens motor nerves. (A, B)** Comparison of the discharge modulation depth (**A**) during sinusoidal head rotation in darkness (0.5 Hz, peak velocity: ±31.4°/s; see peri-stimulus time histograms in Figure 4) and spontaneous firing rate in the absence of head motion (stationary) in darkness (**B**) of the left and right abducens motor nerve in controls (**A_1,_ B**_1_) and one-eared animals (**A_2_, B_2_)**. **(C)** Heat maps of peri-stimulus time histograms depicting the individual activity modulation index (discharge rate/resting rate) during rotation for individual controls (blue) and one-eared (orange) animals (from 12-28 and 14-54 cycles in *n* = 10 and *n* = 15 controls and one-eared animals, respectively; see Figure 4); horizontal rows represent data from individual animals; comparisons of modulation depth and resting rate obtained from controls and one-eared animals for left **(D_1_)** and right **(D_2_)** abducens nerves. Significance levels are indicated by asterisks: * *p* ≤ 0.05, ** *p* ≤ 0.01 (Mann-Whitney *U*-test); ns, no significance (Wilcoxon signed-rank test in **A, B**; Mann-Whitney *U*-test in **D**).

