## Supplementary figures and images for "Strategies of developmental vestibular plasticity after unilateral embryonic ear removal in *Xenopus laevis*"

### Supplemental figures

Supplemental figure 1

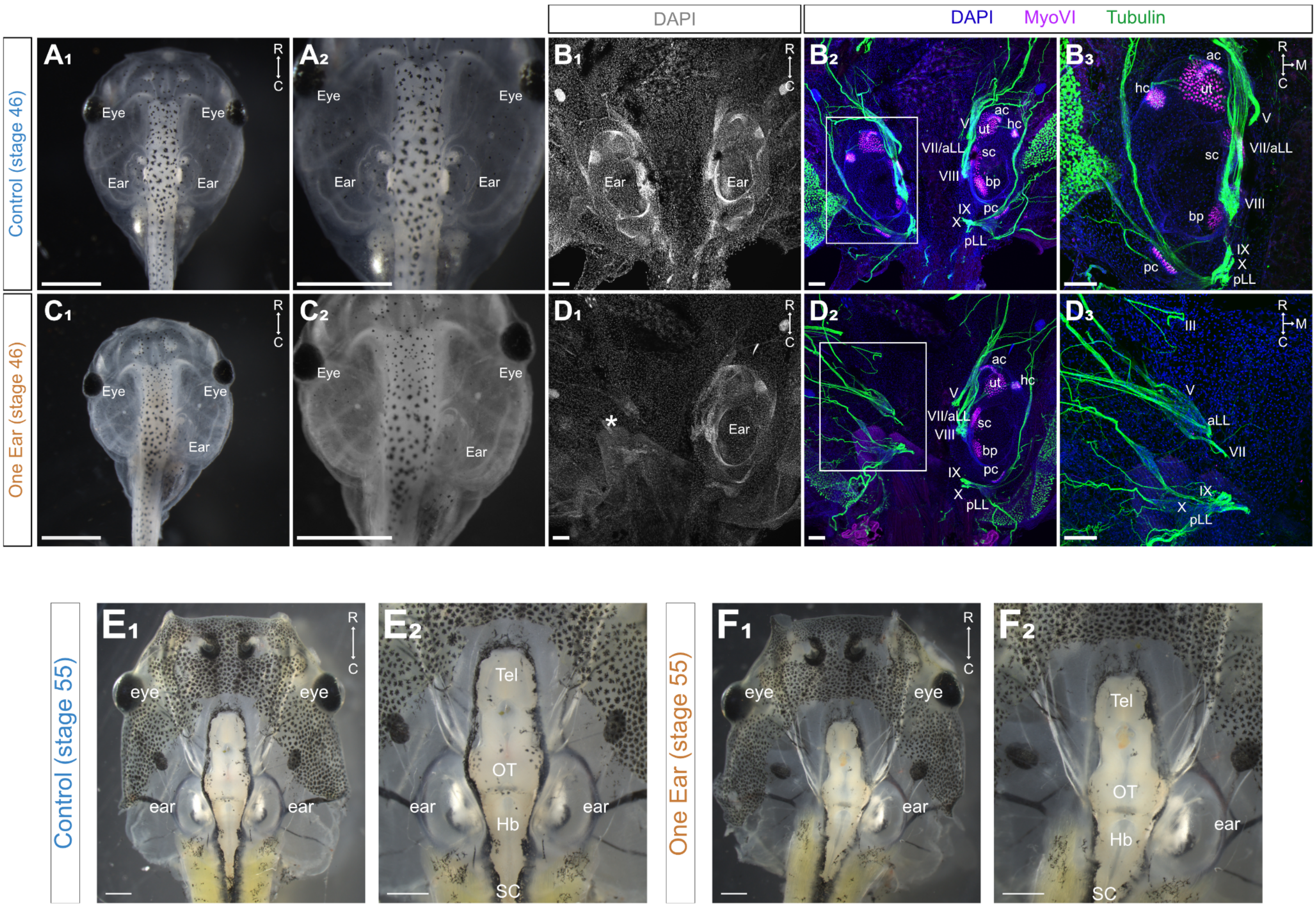

Supplemental figure 2

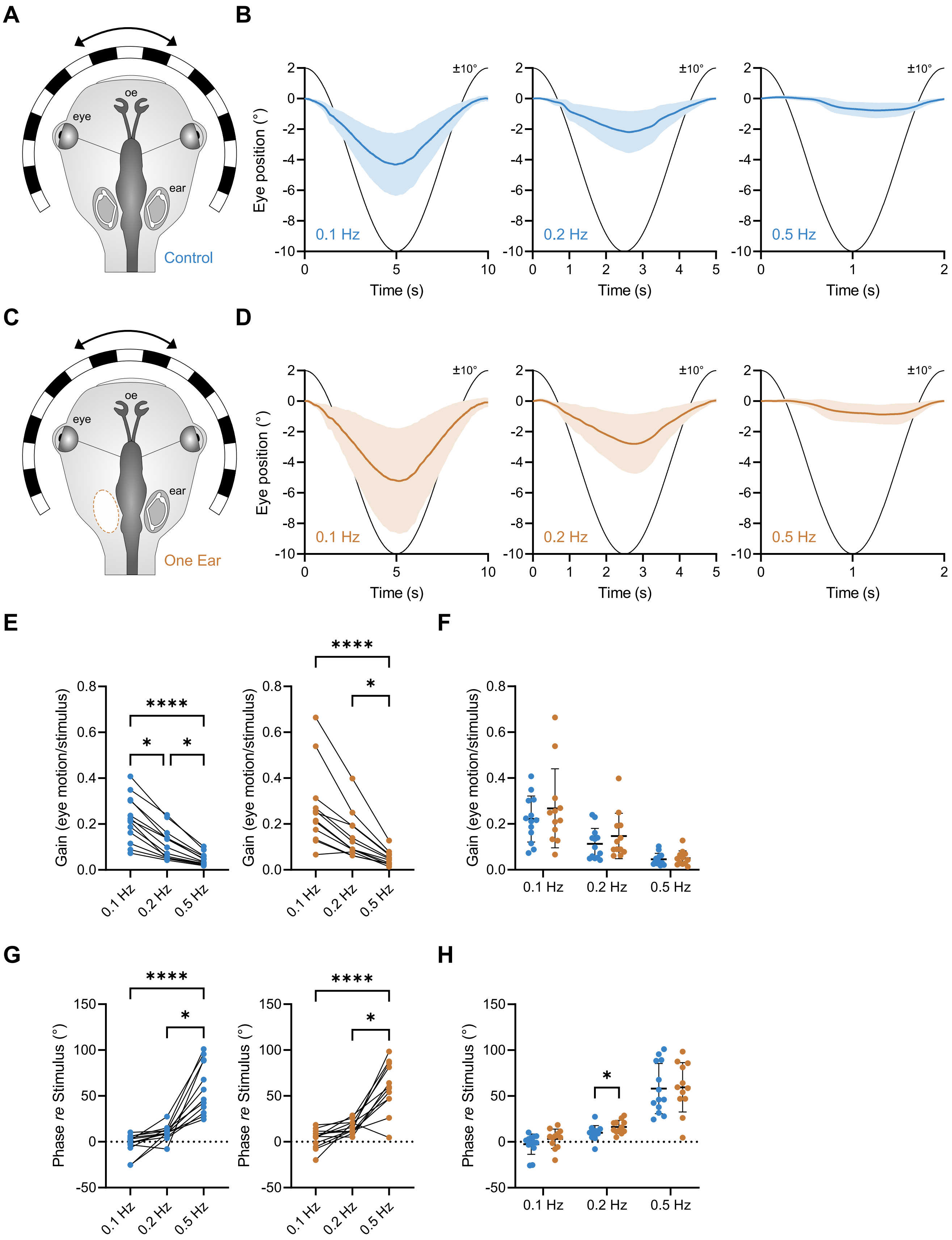

Supplemental figure 3

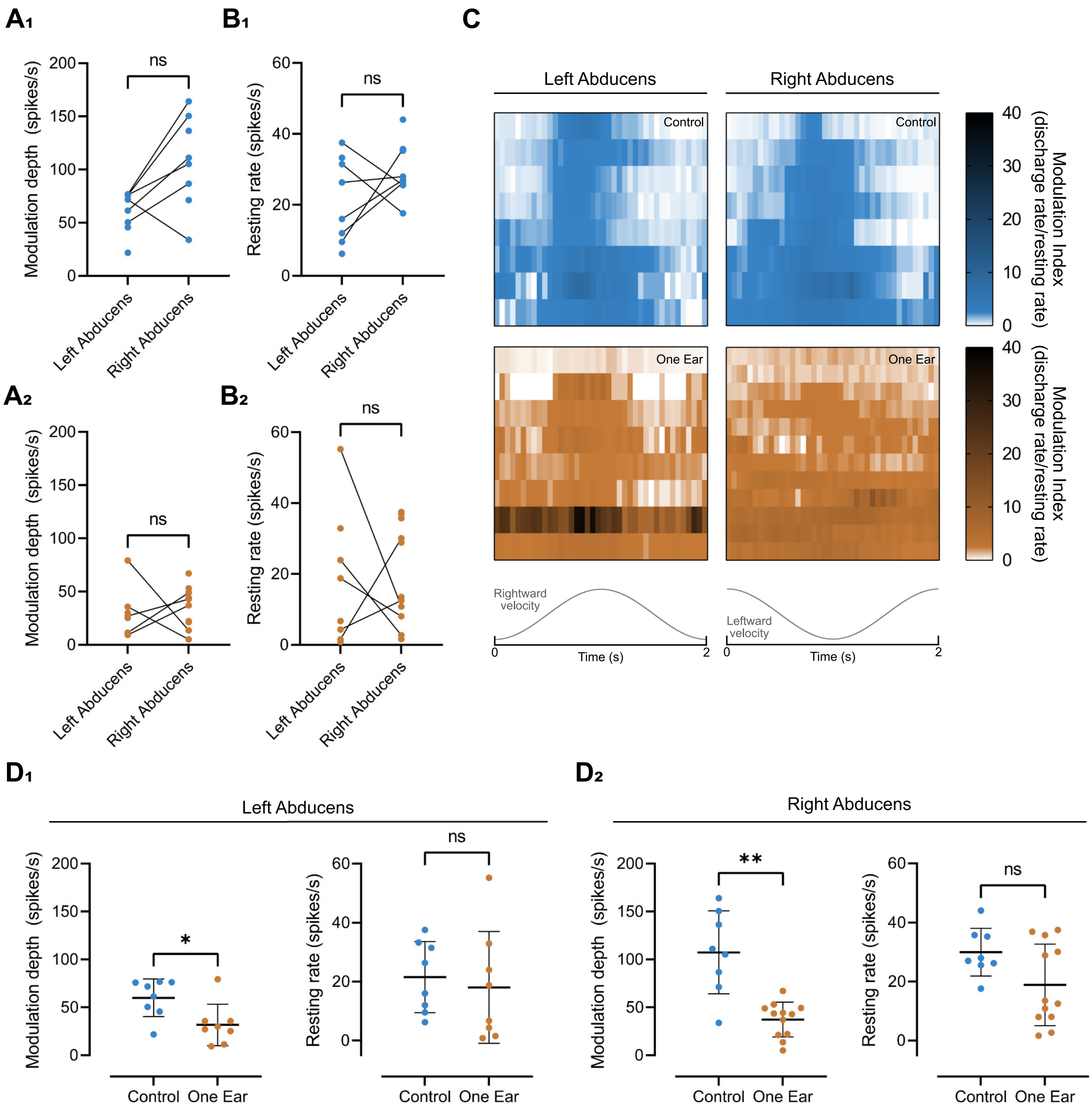
